# Cell-type Specific Ribosomal Tagging Allows for Simultaneous Multi-Tissue Translatomic Sequencing

**DOI:** 10.1101/2025.02.06.636529

**Authors:** Jason Eng, Seock-won Youn, Jan Kitajewski

## Abstract

*In vivo* transcriptomic analysis has advanced significantly with the development of single cell technologies. However, bulk RNA sequencing also continues to provide information on critical signaling pathways and cellular responses. The isolation of mRNA by polysome immunoprecipitation can identify genes undergoing active translation. Unfortunately, the inability to profile multiple cell types from the same sample remains a major downfall of the technique and limits broader analysis of the tissue microenvironment. In this study, we demonstrated the feasibility of immunoprecipitating polysome-associated mRNA from different cell types by strategically expressing differently tagged Rpl22 subunits. Using this technique, we isolated two distinct sets of high quality transcripts from intact B16F10 melanomas, endothelial cells and B16F10 tumors cells, for further molecular analysis.

## Introduction

RNA sequencing techniques have evolved from *en masse* whole tissue to high resolution single cell analysis. As these technologies continue to advance, re-evaluation of methodologies allows for improvements which can facilitate data acquisition while maintaining costs and resources at a manageable level. Recently, single cell sequencing has emerged as the gold standard for transcriptional analysis due to the ability to identify and analyze the genetic response in individual cells^1^. Modifications to this technique, including the development of single nuclei sequencing and ATAC single cell sequencing, have allowed investigators to explore different facets of gene regulation and expression in exquisite detail. However, these approaches have limitations including loss of cytoplasmic transcripts from single nuclei isolation or contamination by signals from dead and stressed cells due to cellular isolation^2^. Moreover, the presence of messenger transcripts does not necessarily correlate with protein production; thus, other modalities to connect gene signatures with protein production can provide vital information regarding cellular activity.

The RiboTag mouse system is an *in vivo* transgenic model developed by McKnight *et al* for the isolation of mRNA by ribosomal immunoprecipitation^3^. In these mice, *loxP* sites flank the C-terminal exon and stop codon of the native ribosomal protein L22 (*Rpl22*). An identical copy of the exon with multiple HA-tag repeats is inserted downstream of the terminal site. When crossed with a Cre recombinase driven by a tissue specific promoter, the native C-terminal exon is replaced in selective cell types with the HA-tagged exon sequence. The resulting Rpl22-HA subunit incorporates into ribosomes and participates in the translation of mRNA into peptide sequences^4^. Compared with other methodologies, RiboTag offers several advantages. To generate tissue specific sequences in single cell RNA sequencing, cell isolation by various techniques such as flow cytometry or magnetic bead sorting is needed prior to harvesting RNA. However, studies have shown that cell sorting can cause upregulation of stress response genes which may skew signaling pathway analysis^5^. In contrast, mRNA isolated using the RiboTag system can be performed from frozen tissues without the need for prolonged digestion and manipulation of individual cells, thereby eliminating cell stress and offering a more relevant picture of the genetic response.

Here, we adapted the *in vivo* isolation of polysomes using the RiboTag mouse system to immunoprecipitate translating mRNA from different cell types. Our studies build upon techniques for polysome immunoprecipitation by using murine tumor cells transfected to express uniquely tagged ribosomal subunits. Once implanted into a RiboTag mouse, the translating mRNA from both the tumor cells and a surrounding cell type of interest can be simultaneously captured and separately interrogated. To demonstrate the feasibility of this technique, we isolated tumor and endothelial specific transcripts from samples without the need for purification of live cells or intact organelle structures. Our findings show that this method produces a clean translatomic signature for at least two cell types, while also limiting confounding stress and cellular disruption which could impact downstream pathway analyses.

## Materials and Methods

### Cell Lines

B16F10 melanoma tumor cells and human embryonic kidney cells (HEK293T) were acquired from ATCC. These cells were cultured in high glucose DMEM (Gibco) supplemented with 10% fetal bovine serum (Avantor) and penicillin-streptomycin (Gibco) at 37^°^C at 5% CO_2_. The cell line was tested monthly for mycoplasma contamination using the MycoAlert kit (Lonza).

### Subcloning for Ribosomal Protein L22

Total RNA was isolated with Trizol (Invitrogen) from normal murine lung tissue. Mouse cDNA was synthesized with High Capacity cDNA reverse transcription kit (Applied Biosystems). The murine ribosomal protein L22 (*Rpl22*) was amplified with Phusion Taq (NEB) using mRpl22 primers and inserted into pCCL-E2A-mCherry-T2A-Puromycin vector. To insert 3xFlags, a double stranded sequence of 3xFlag was synthesized and ligated into the C-terminal pCCL-Rpl22-mCherry-Puro cut with XbaI (NEB) and SalI (NEB). The primer sequences are shown in Table 1.

**Table 1.**
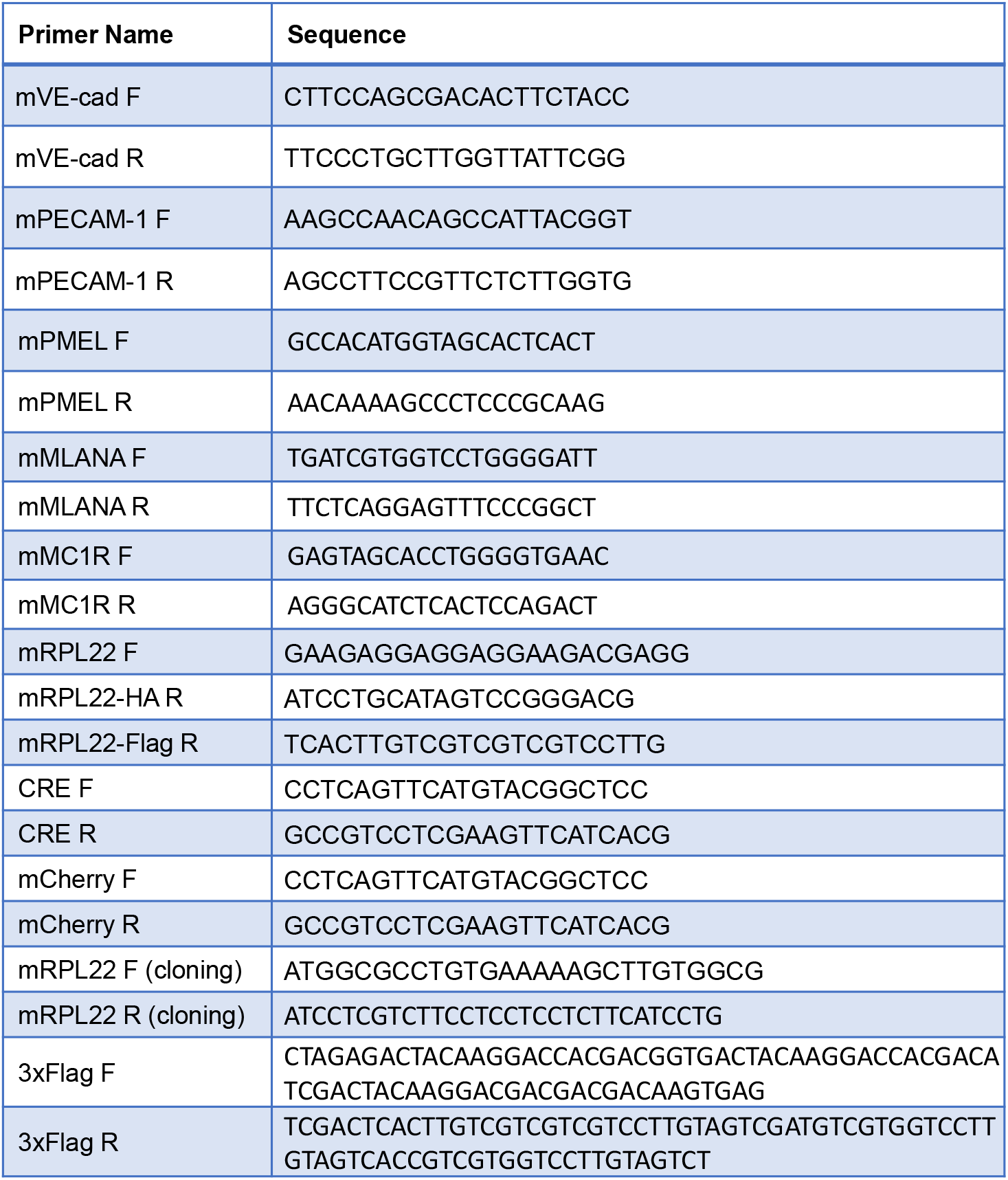
Sequences of murine qPCR primers and cloning primers.

### Lentivirus Synthesis and Transduction

HEK293T cells were cultured at 70% confluence in DMEM with 10% FBS without antibiotics. 1mg/ml of polyethyleneimine (Polysciences) was used to transfect lentiviral packaging vectors (pVSVg, pMDLg/pRRE, pRSV/REV) and target genes (pCCL-Luciferase-E2A-mCherry-T2A-Puromycin or pCCL-Rpl22-3xFlag-E2A-mCherry-T2A-Puromycin). After an overnight incubation, the transfection media was replaced with fresh DMEM with non-essential amino acid (Gibco). The lentiviral media was collected and then passed through a 0.45μm filter (Thermo Scientific). For transduction, the B16F10 cells were incubated overnight with purified lentivirus media and 8μg/ml of polybrene (Sigma). Transfection efficiency was determined by the ratio of mCherry expressing cells over the total cell number at Day 3 post-transduction. To enrich the population, 1μg/ml of puromycin (Gibco) was added to the culture for 3 days to select for stably transfected cells.

### Western Blot

The lentivirus transduced B16F10 cells were lysed with RIPA buffer: 50mM Tris pH 8.0 (Invitrogen), 150mM NaCl (Research Products), 1% Triton X-100 (Fisher Sci), 0.5% sodium deoxycholate (Sigma), 0.1% SDS (Fisher Sci), 2mM EDTA (EMD Millipore) with Halt protease inhibitor cocktail (Invitrogen). 40μg of total protein was separated by 12% SDS-PAGE and transferred onto 0.22μm PVDF membrane (Biorad). Anti-Flag (Cell signaling), anti-mCherry (Abcam), and anti-a-tubulin (Sigma) were used to analyze gene expression.

### RiboTag Mice Maintenance

All mice were housed in the animal research facility at the University of Illinois Chicago in accordance with approved IACUC protocols. RiboTag mice were acquired from the McKnight group, while Cdh5-CRE-ER^T2^ mice were provided by Ralf Adams^6^. Homozygous RiboTag with Cdh5-CRE-ER^T2^ mice were maintained for further experiments.

To induce recombination, tamoxifen (Sigma) was dissolved into corn oil (Sigma) at 20 mg/mL. 3 week old male Cdh5-CRE-ER^T2^/RiboTag^+/+^ mice were injected with the tamoxifen solution at 75mg/kg intraperitoneally for 5 days. Afterward, the mice were allowed to mature until 8 weeks of age.

### B16F10 RPL22-FLAG Tumor Implantation

A total of 5×10^5^ B16F10 RPL22-FLAG tumor cells were subcutaneously implanted into both the left and right posterior flanks of 8 week old tamoxifen induced, Cdh5-CRE-ER^T2^/RiboTag mice. After 2 weeks, the mice were humanely euthanized per IACUC guidelines. The right flank tumors were quickly resected and placed into optimal cutting temperature (OCT) (Fischer Sci) solution for histology, while the left flank tumors were snap frozen for polysome isolation.

### Immunofluorescence Staining of Tumor Sections

The tumor blocks were sectioned at 8μm thickness on to Superfrost Plus slides (Fisher) using a Epredia™ Microm HM525 NX Cryostat (Leica). First, the sections were warmed to room temperature and air dried. They were rinsed once in 1x PBS to remove the OCT and then fixed for 7min at -20^°^C with methanol (Fisher Sci). After three rinses in PBS with 0.5% Tween-20 (Sigma) to remove residual methanol, the sections were blocked with 5% normal donkey serum (Vector Lab) in PBS-T at room temperature for 30mins. Antibodies against HA (Abcam) 1:500, FLAG (Biolegend) 1:200, and CD31 (BD Biosciences) 1:500 in blocking buffer were incubated on the sections overnight at 4^°^C. The following day, the sections were washed with PBS-T and incubated for an hour at room temperature with goat anti-rat Alexa 488 (Invitrogen) or goat anti-rabbit Alexa 647 (Invitrogen) 1:1000 in PBS-T. The slides were again washed with PBS-T and then mounted with Vectashield Mounting Media with DAPI (Vector Lab). Images were obtained with an AxioImager (Zeiss) at 10x magnification and channels were merged using ImageJ software.

### Dual RiboTag Immunoprecipitation and Real-time PCR Confirmation

A 50μL aliquot of Dynabeads A (Thermofisher) was washed twice with wash buffer: 50mM Tris pH 7.5 (Invitrogen), 250mM NaCl, 15mM MgCl_2_ (Invitrogen), 100μg/ml of Cycloheximide (Sigma), 0.5% Triton X-100, DNAse/RNAse Free H_2_O (Research Products). Anti-HA (Abcam) or Anti-Flag (Biolegend) antibodies were conjugated to magnetic beads and washed twice with wash buffer. The snap frozen tumors were first homogenized by mortar and pestle in liquid nitrogen, immediately followed by lysis in a glass Dounce homogenizer with 1:10 tissue mass to lysis buffer volume: 50mM Tris pH 7.5, 100mM KCl (Invitrogen), 15mM MgCl_2_, 100μg/mL of Cycloheximide, 0.5% Triton X-100, 1 mM DTT (Sigma), 7μL/mL Protector RNase inhibitor (Sigma), 12μL/mL Turbo DNase (Invitrogen), Halt protease inhibitor cocktail (Invitrogen). The tissue was homogenized with 10 strokes each of pestle A and B. The lysate was centrifuged at 10,000xG at 4°C for 10min to remove debris. 800μL of supernatant was carefully collected and incubated with 50μL of antibody/Dynabead mixture for 2 hours rocking on ice. After the incubation, the beads were collected with magnetic columns and washed twice with bead wash buffer: 50 mM Tris pH 7.5, 100 mM KCl, 15 mM MgCl_2_, 100μg/ml of Cycloheximide, 1% Triton X-100, 1mM DTT, 1μL/mL Protector RNase inhibitor, 8μL/mL Turbo DNase, Halt protease inhibitor cocktail. mRNA was isolated from the beads with an RNeasyPlus kit (Qiagen). The RNA integrity number (RIN) score was measured with the LifeTechnologies Tapestation. The isolated mRNA (1μg) was reverse transcribed to cDNA with a High Capacity cDNA reverse transcription kit (Applied Biosystems). To validate cell specific isolation, endogenous genes for endothelial markers (*Pecam1, Cdh5*) ^7^ and melanoma markers (*Pmel, Mlana, Mc1r*) ^8,9^ and exogenous genes for endothelial cells (*Rpl22*-*HA, Cre*) and melanoma cells (*Rpl22-Flag, mCherry*) were analyzed by real-time PCR with primers (Table 1). The reaction was performed using Fast SYBR green master mix (Applied Biosystems) with ABI Viia7 machine (Applied Biosystems). The DDCT method was used to calculate differences in the gene expression.

### Statistics

For experiments, unless otherwise noted, t-test with unpaired analysis was performed on all quantified data to determine significant differences between groups using GraphPad Prism 9. P-values less than 0.05 were considered statistically significant. p-values < 0.05 are flagged with one star (^*^), p-values < 0.01 with two stars (^**^), p-values < 0.001 with three stars (^***^), and p-values < 0.0001 with four stars (^****^). Unless otherwise noted, experiments were repeated at least three times.

## Results

### Development of the Rpl22-FLAG Construct

To isolate translating mRNA, we first devised a strategy to tag the RPL22 subunits with unique epitopes to facilitate immunoprecipitation. We employed the RiboTag mouse model, which expresses HA-tagged RPL22 subunits, to harvest transcripts from tissues *in situ*. For a second isolation method, we generated a construct containing the *Rpl22* transcript fused with a 3xFLAG tag and an mCherry reporter under the control of the PGK promoter, referred to as RPL22-FLAG. The resulting plasmid was packaged into a lentiviral vector for efficient cellular infection. To maximize the extraction of ribosomal subunits, a puromycin selection marker was also incorporated into the plasmid to positively select for cells stably expressing the RPL22-FLAG protein.

After successful production of the lentivirus, we infected B16F10 murine melanoma cells and selected the stably expressing cells with puromycin. The viable cells were injected subcutaneously into the posterior lateral flank of tamoxifen-induced Cdh5-CRE-ER^T2^/RiboTag^+/+^ mice. Once tumors had grown to sufficient size, the mice were humanly euthanized per IACUC protocol, and the tissue was rapidly snap frozen or placed in OCT (Figure 1).

**Figure 1.**
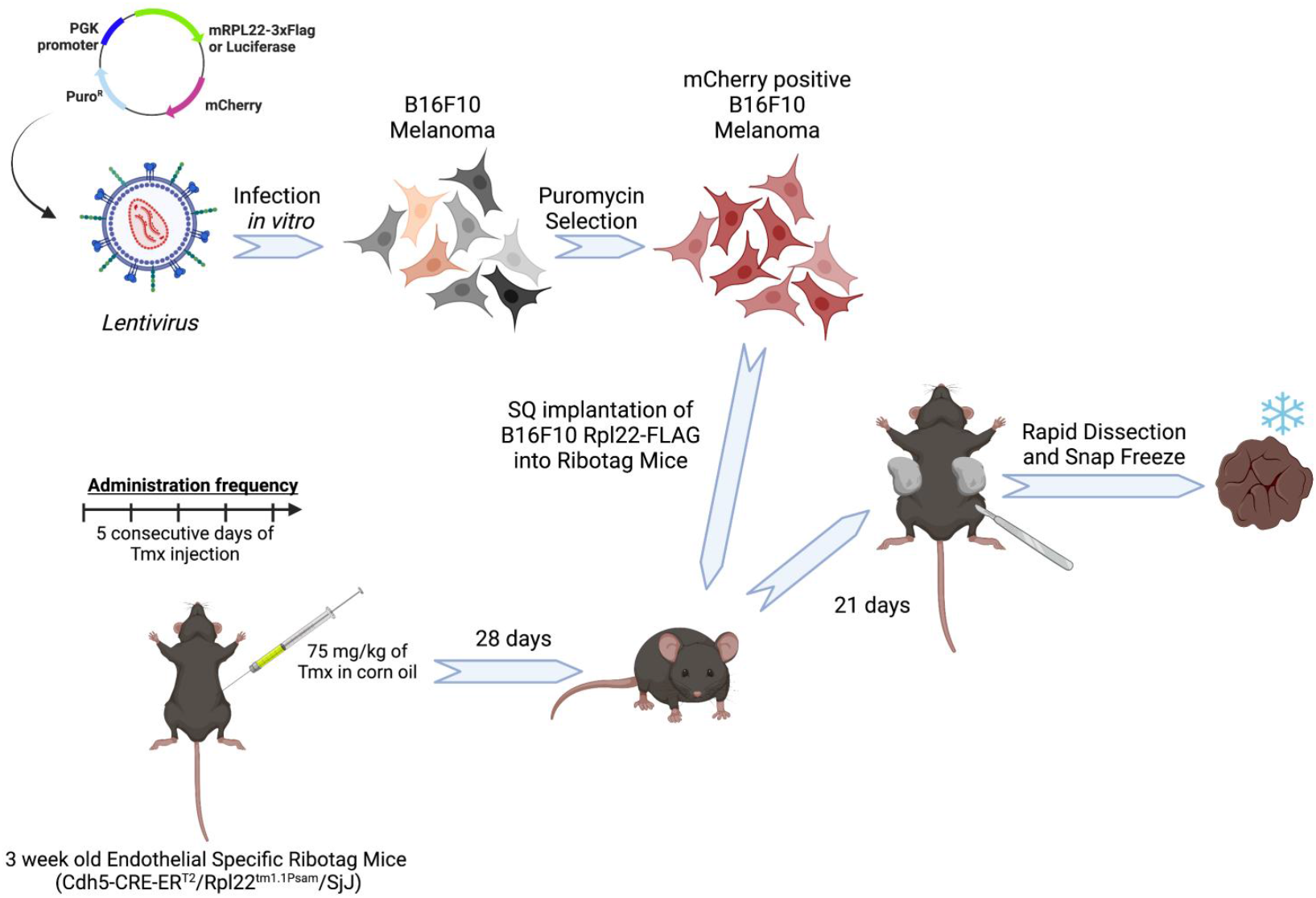
Schematic illustrating the development of the lentivirus used to introduce the RPL22-FLAG construct into B16F10 melanoma cells, followed by implantation of the tumors into C57BL/6j mice.

### Validation of Rpl22-HA and Rpl22-FLAG Expression

Expression of RPL22-FLAG was first validated by western blot analysis comparing the expression of the mCherry tag and the FLAG tag in the parental wildtype tumor cell line as well as in a separate B16F10 line transfected with a construct that substituted the RPL22-FLAG sequence for luciferase (Figure 2A). Additionally, fresh frozen tissue sections were stained for both FLAG and HA expression for further confirmation. As anticipated, HA and the FLAG-tag expression did not overlap indicating successful labeling of two separate cell populations within the tumor (Figure 2B). To further validate the HA expression was indeed localized to the endothelial cells, CD31 and HA co-staining was performed. The overlapping signals demonstrate tumor blood vessel specific expression of HA (Figure 2C).

**Figure 2.**
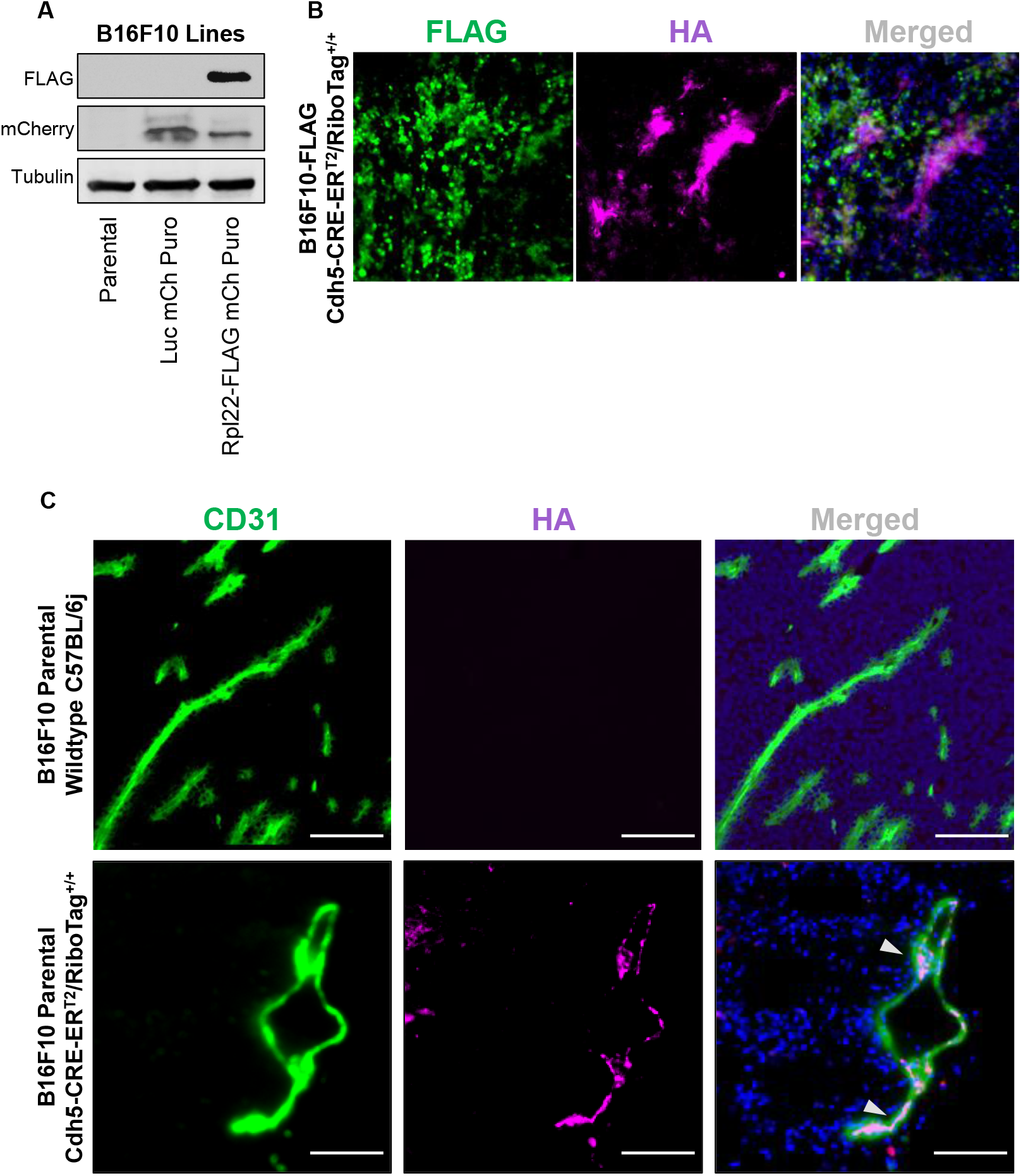
Validation of the RPL22-FLAG tag. A) Western blot analysis of the B16F10 cell line including the non-transfected parental line, a control plasmid infected line containing luciferase, and the RPL22-FLAG containing construct. Expression of the FLAG-tag and mCherry-tag were validated prior to implantation into mice. B) FLAG-tag and HA-tag expression assessed by immunofluorescence in B16F10-FLAG tumors grown in tamoxifen-induced endothelial specific RiboTag mice. C) Validation of endothelial specific HA-tag expression in B16F10-FLAG tumors implanted into tamoxifen-induced RiboTag mice compared with B16F10 parental cells implanted in wildtype C57BL/6j mice.

### Optimization of Rpl22 Pulldowns

Efficient immunoprecipitation of RPL22 subunits is crucial for quality mRNA purification. To ensure sufficient yield, tissues were first finely pulverized in liquid nitrogen with cleaned mortars and pestles. The ground tissue was quickly transferred into lysis buffer and placed into Dounce homogenizers. Immunoprecipitation with anti-HA antibodies and anti-FLAG antibodies from the lysate was performed and the resulting mRNA was isolated (Figure 3A). RIN scoring of the isolated RNA revealed adequate quality material for further use in the majority of molecular studies (Figure 3B).

**Figure 3.**
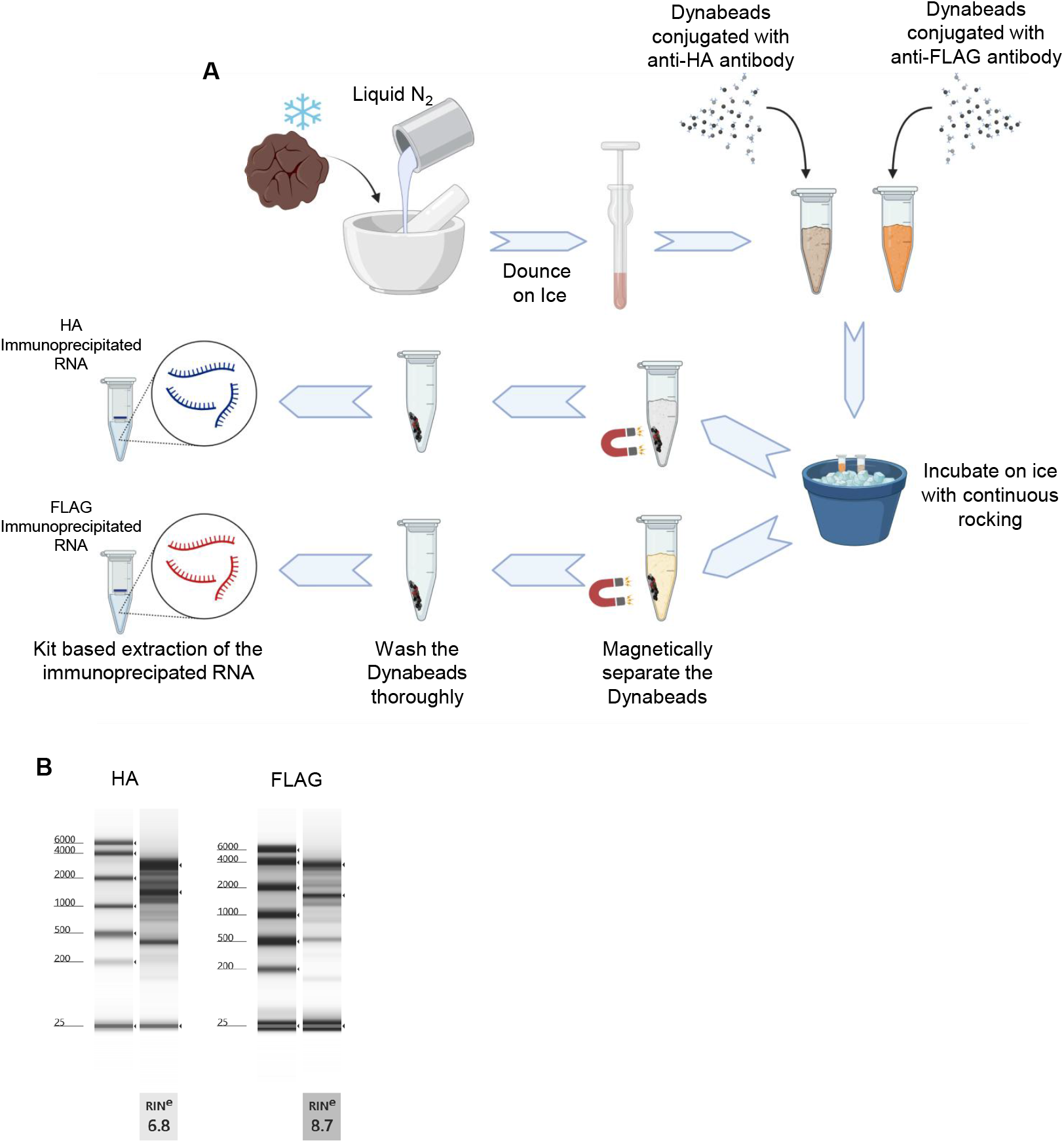
Processing and quality control of the dual RiboTag. A) Schematic illustrating the isolation of ribosomal associated mRNA from B16F10-FLAG tumors implanted in tamoxifen-induced endothelial specific RiboTag mice. (B)RIN scores showing the quality of RNA isolated by immunoprecipitation with RPL22-HA or RPL22-FLAG.

### Real-time PCR validation for Tissue Specific Transcripts

To validate the tissue specific isolation of mRNA from tumor cells by FLAG immunoprecipitation and from endothelial cells by HA immunoprecipitation, we performed real-time PCR analysis of melanoma and vascular markers. The expression of endothelial markers, *Pecam1* and *Cdh5*, was significantly enriched in the HA-IP fraction compared with the FLAG-IP fraction (Figure 4A). Conversely, the expression of melanoma associated genes, *Mlana* and *Mc1r*, were significantly enriched in the FLAG-IP fraction compared with the HA pulldown (Figure 4B). Interestingly, while the tumor antigen, gp100 or *Pmel*, was also increased in the FLAG pulldown, it was not statistically significant due to the variability in the expression of the gene.

**Figure 4.**
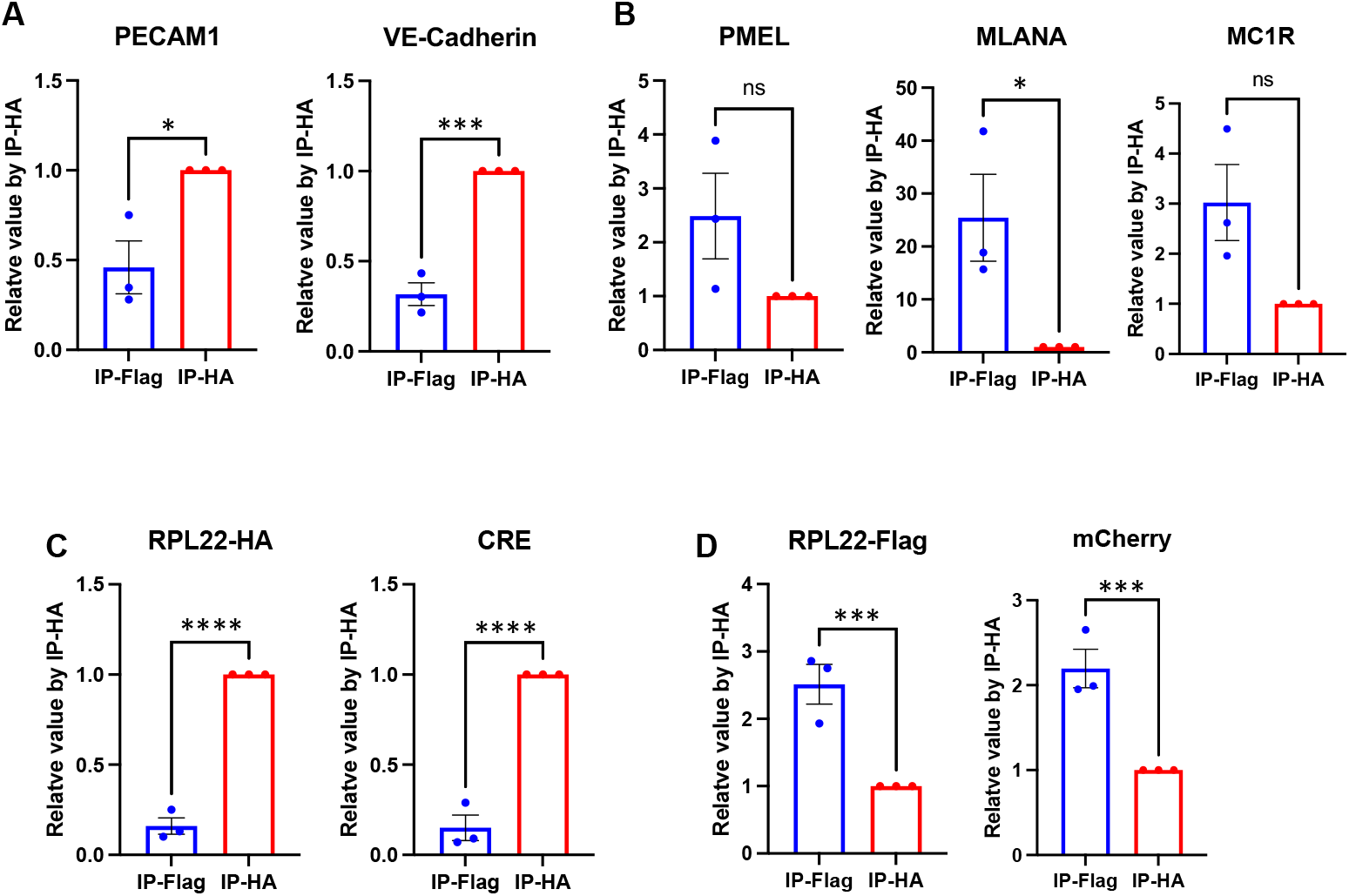
Validation of the dual RiboTag System. RT-PCR analysis of A) endothelial associated genes, *Pecam1* and *VE-Cadherin/Cdh5*, and B) melanoma associated genes, *Pmel, Mlana, and Mc1r*, from HA-IP and FLAG-IP. (C)Additional validation of tag specific immunoprecipitation using RT-PCR analysis of the C) exogenous endothelial specific HA-tag and Cre recombinase gene, as well as the D) exogenous tumor associated FLAG-tag and mCherry-tag isolated fractions. *n=3* tumors/group.

Additionally, we further verified the specificity of the immunoprecipitated ribosomal captured RNA by assessing the expression of the exogenous HA and FLAG-tags. The endothelial-expressed RPL22-HA was significantly enriched in the HA immunoprecipitation (Figure 4C), while FLAG transcripts were notably expressed in the FLAG-IP fraction associated with melanoma genes (Figure 4D). Reassuringly, the levels of the tag transcripts in the alternate immunoprecipitated fractions, i.e. FLAG expression in the HA-IP, was low indicating minimal contamination from non-specific pulldown.

## Discussion

Gene analysis relies on capturing high quality transcripts while also minimizing external factors that alter the readout. Single cell RNA sequencing has expanded the information available for biomedical research; however, limitations of the technique, including the potential introduction of artifactual reads due to cell isolation-induced stress, can distort findings. To address this issue, techniques which limit or avoid individual cell disaggregation steps may provide a truer snapshot of intrinsic cell responses. Recently, the ability to isolate RNA from frozen or even fixed cells has allowed for examination of previously difficult to dissociate tissues that would typically require aggressive enzymatic and mechanical disruption. These methods still have certain drawbacks. Namely, the isolation of transcript rich organelles, such as the nucleus, can lose cytoplasmic RNA and decrease the yield of genes that are not undergoing active transcription. In addition, the presence of transcripts within the nucleus does not always correlate to translated proteins which may inaccurately place greater significance on nuclear mRNA. Therefore, the RiboTag system provides additional data to bridge the gaps from transcribed messengers to protein.

The translatome derived from the immunoprecipitated RNA allows for a deeper understanding of signaling pathways critically regulated during a cellular response. Our work presented here highlights the possibility of expanding the utility of the RiboTag system. Previous studies were limited by the availability of a single tag under the control of one selected promoter. The HA-tag proved to be an opportune epitope due to its small size, which minimized protein folding interference, and the availability of high affinity antibodies, which facilitated immunoprecipitation. We ultimately employed the FLAG-tag for similar reasons but enhanced the epitope availability by utilizing three FLAG-tags in tandem. To ensure adequate RNA isolation, we also chose to perform the FLAG-IP from the most abundant cell population, while the Cdh5-Cre-ER^T2^ driven endothelial specific HA-IP was utilized for the endothelial population in the tumor. We believe that this strategy allowed for optimal mRNA yield from all the desired tissues. Gregory *et al* demonstrated the possibility of multi-cell type polysome isolation from an *in vitro* mixed cell population with different RPL22 expressing tags^10^. Interestingly, they excluded the FLAG tag because preliminary validation studies with co-IP were unsuccessful at pulling down the RPL0P0 subunit with RPL22-FLAG. However, these findings would not necessarily affect the RiboTag assay since RPL22 does not directly interact with RPLP0 within 60S ribosomal complex. Furthermore, RPL22 alone can bind mRNA while RPLP0 does not^11^. Based on our results, the RPL22-FLAG can precipitate tissue specific mRNA from samples demonstrating that it is indeed functional in our system.

Even though the cancer cell population comprised a large portion of the tumor, it was evident by our quality analysis, the yield of mRNA was low. This was likely due to multiple factors including low tumor cell lysis, decreased FLAG-tag expression over time and suboptimal solubilization of the polysomes leading to low release of the mRNA from the ribosome complex. While retention of the RPL22-FLAG expression may prove difficult to control, more complete cell lysis along with better protein and nucleic acid solubilization, such as with Trizol reagent, may improve the yield^12^. Nevertheless, this method was able to extract mRNA of high enough quality for molecular analysis including qPCR and likely RNA sequencing. Overall, our findings here highlight a new advancement for the RiboTag technique that could help provide additional information that may be overlooked or obscured by current single cell methods.

## Future Perspectives

We anticipate that with the development of more advanced protocols for the delivery of recombinant DNA, that utility of a multi-tag RiboTag systems will expand. Currently, the ability to introduce specific *Rpl22* transcripts is limited by the mode of delivery, such as viral and nanoparticle based systems. As the capability to deliver cell type specific transcripts improves, the potential to introduce multiple tags will also provide an opportunity to isolate different cell type translatomic signatures from a single segment of tissue. This data would help guide understanding of which genes play a crucial role in disease initiation, progression, and therapeutic response.

## Executive Summary

Translatomic analysis provides additional data on top of conventional transcriptomic techniques, but it is limited in its ability to effectively interrogate multiple cell types *in vivo*.

We developed a strategy to isolate ribosomes and the associated mRNA from a single tumor using the RiboTag (RPL22-HA) murine mouse model and a tumor line that was transduced by lentivirus with RPL22-FLAG.

We captured mRNA from both endothelial cells and B16F10 melanoma cells within the same tumor and demonstrated tissue-specific gene expression using qPCR analysis.

Additional complexity for this technique can be incorporated by employing further unique tags for ribosomal precipitation in order to investigate other cell-types and supplement transcriptomic analysis.

## Financial Disclosures/Acknowledgements

The authors have no relevant financial disclosures.

## Ethical Conduct of Research

All animal studies were conducted in accordance with guidelines put forth by the IACUC at the University of Illinois Chicago.

